# Calcium-induced environmental adaptability of the blood protein vitronectin

**DOI:** 10.1101/2022.03.10.483860

**Authors:** Ye Tian, Kyungsoo Shin, Alexander E. Aleshin, Wonpil Im, Francesca M. Marassi

**Affiliations:** Sanford Burnham Prebys Medical Discovery Institute, 10901 North Torrey Pines Road, La Jolla CA, 92037; Departments of Biological Sciences, Chemistry, and Bioengineering, Lehigh University, PA 18015, USA

**Author notes:** **Correspondence:** Francesca M. Marassi, Sanford Burnham Prebys Medical Discovery Institute, 10901 North Torrey Pines Road, La Jolla CA, 92037, USA.

**Keywords:** Vitronectin, structure, calcium, calcification, blood, shear stress, macular degeneration

## Abstract

The adaptability of proteins to their work environments is fundamental for cellular life. Here we describe how the hemopexin-like (HX) domain of the multifunctional blood glycoprotein vitronectin (Vn) binds Ca^2+^ to adapt to excursions of temperature and shear stress. Using X-ray crystallography and molecular dynamics (MD) simulations, nuclear magnetic resonance (NMR) and differential scanning calorimetry (DSF), we describe how Ca^2+^ and its flexible hydration shell enable the protein to perform conformational changes, that relay beyond the Ca^2+^ binding site to alter the number of polar contacts and confer conformational stability. By means of mutagenesis, we identify key residues that cooperate with Ca^2+^ to promote protein stability, and we show that Ca^2+^ association confers protection against shear stress, a property that provides conformational advantage for proteins that circulate in the vasculature like Vn. The data reveal a mechanism of adaptation.

**Significance Statement:** The protein vitronectin (Vn) plays important roles in cell adhesion and migration, bone remodeling and immunity. It circulates in blood, but it is also found in the extracellular matrix, and it accumulates with plaques associated with age-related macular degeneration, Alzheimer’s disease, atherosclerosis and other degenerative disorders. Vn is a calcium-binding protein, and here, we show that calcium helps Vn alter its structure in response to diverse environmental conditions. The results shed light on the way in which Vn adapts to its surroundings. This structural knowledge is important for the development of diagnostic, preventive or therapeutic approaches.

---

Proteins in the vasculature are particularly adept at functioning in a diverse range of chemical and physical conditions. Vitronectin (Vn) circulates in blood at a concentration of 0.2–0.5 mg/ml and accounts for 0.1–0.5% of plasma protein (1-3), but it is also a component of high-density lipoprotein (4, 5), and associates with the extracellular matrix through collagen-binding and heparin-binding domains (6). Vn interacts with multiple ligands to regulate diverse physiological processes: It employs its Arg-Gly-Asp sequence to bind integrin receptors and promote cell adhesion, spreading and migration (7); regulates hemostasis through the interactions of its somatomedin B domain with type 1 plasminogen activator inhibitor (8); controls complement-mediated pore formation and cell lysis through the interactions of its C-terminal domain with the C5b-C9 components of the membrane attack complex(9); and is recruited by many pathogens to acquire protection from complement-mediated lysis (10).

While Vn is synthesized primarily in the liver, it is also expressed in the retina (11-13), brain (14), and vascular smooth muscle cells (15). Moreover, Vn expression is upregulated in a number of diseases, including fibrotic tissues (16), atherosclerosis (17), Alzheimer disease (AD) (18-20), and age-related macular degeneration (AMD) (20, 21). In these settings, Vn is often associated with calcified protein-lipid deposits that accumulate with disease progression (22), but the specific role of Vn in calcified deposit formation is not well understood.

Previously, we determined the three-dimensional structure of the hemopexin-like (HX) domain of Vn (23). We showed that it specifically binds both soluble Ca^2+^ and mineralized hydroxyapatite [Ca_10_(PO_4_)_6_(OH)_2_] – the major component of pathological calcified protein-lipid deposits – acquiring appreciable thermodynamic stability in the process (24). Here we describe the structural mechanism for this Ca^2+^-dependent effect, and show that it also confers Vn HX with mechanical stability against high shear stress, such as that of the vasculature. Like other blood proteins, Vn has evolved to function in a diverse range of chemical and physical environments. We show that Ca^2+^ association with key Vn residues enables the protein to adapt to environmental extremes of temperature and shear stress by stabilizing multiple protein states with subtly different conformations and levels of hydration. These results are important for understanding the roles of Vn in health and disease, and also illustrate a fundamental mechanism of environmental adaptation.

## Results and Discussion

### Structure of Ca^2+^-bound Vn HX domain

To resolve the Ca^2+^ binding site of the HX domain, we determined two structures, each from a crystal treated with 1 mM CaCl_2_ (Vn-Ca_1_) or 100 mM CaCl_2_ (Vn-Ca_2_). The structures refined to a resolution of 2.0 Å and 1.7 Å, respectively (Table S1). They have the same crystallographic space group as the previously determined (23) Na^+^-bound structure (Vn-Na_1_), but Vn-Ca_2_ has different unit cell parameters and different crystal packing. Soaking the Vn-Ca_2_ crystal with EDTA yielded a second Na^+^-bound and Ca^2+^-free structure (Vn-Na_2_), with the same space group and unit cell parameters as Vn-Ca_2_ and a resolution of 2.0 Å. In all structures, the crystallographic asymmetric unit contains two copies of the protein molecule. The Vn-Ca_2_ and Vn-Na_2_ structures are representative of the differences between Ca^2+^ and Na^+^ bound states (Fig. 1).

**Fig. 1.**
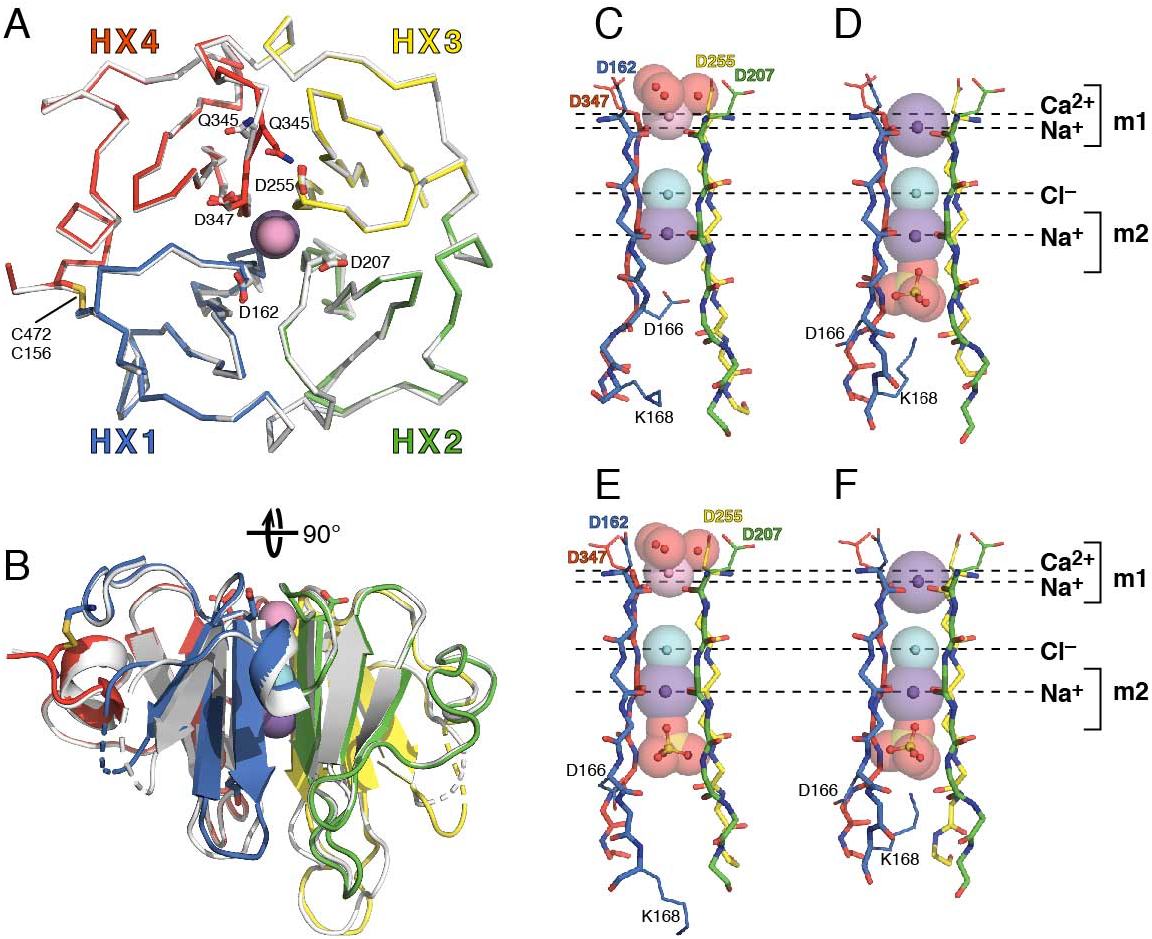
Structures of Ca^2+^-bound Vn. Colors denote structural repeats HX1 (blue), HX2 (green), HX3 (yellow), and HX4 (red). Spheres denote Na^+^ (indigo), Ca^2+^ (pink), Cl^-^ (cyan), and O (red) or S (yellow) from water or bound SO_4_^2^. Broken loops denote gaps in the protein chain due to missing electron density. Sticks denote key sidechains, disulfide linked Cys, and the backbone of the central channel. **(A, B)** Top and side views of Vn-Ca_2_ (color) and aligned Vn-Na_2_ (gray). **(C-F)** Backbone representations of the channel structures of Vn-Ca_2_ (C, E) and Vn-Na_2_ (D, F). Molecules A (C, D) and B (E, F) of the asymmetric unit are shown for each structure. Dashed horizontal lines mark the positions of Ca^2+^ and Na^+^ (at sites m1 and m2) and Cl^−^ ions.

Ca^2+^-bound Vn HX maintains a four-bladed β/α propeller, where each of the four HX repeat sequences corresponds to one propeller blade (β1-β2-β3-α) and the termini are linked by a C156-C472 disulfide bond (Fig. 1A, B). The propeller top – defined as the β1 N-termini – forms a smooth surface while loops of variable lengths protrude from the bottom, including two flexible regions, spanning residues 283-324 and 354-435, which were deleted to facilitate refolding and crystallization (23). The propeller bottom has overall higher B factors and some missing electron density for the more flexible regions.

The four β1 strands form a channel (Fig. 1C-F; Fig. S1) that occludes a NaCl molecule, with Na^+^ coordinated by the backbone carbonyl oxygens of β1 residues F164, A209, A257 and A349 (metal binding site m2) and Cl^−^ coordinated by the backbone amide hydrogens of A163, A207, A256 and A348.

Four Asp sidechains (D162, D207, D255, D347) extend from the top rim of the channel to form an electronegative pole (23, 24). The backbone carbonyl oxygens of these rim-Asp coordinate a Ca^2+^ or Na^+^ ion (metal binding site m1). The Ca^2+^ coordination sphere is completed by three (Vn-Ca_2_) to four (Vn-Ca_1_) water molecules. Several of the Ca^2+^-bound waters are within 3.2 Å of the rim-Asp carboxyl oxygens and establish a water-mediated H-bond network involving the rim-Asp sidechains, water and Ca^2+^. By contrast, Na^+^ is either not hydrated (Vn-Na_2_) or bound to a single water molecule (Vn-Na_1_).

Ca^2+^ ligation is supported by agreement of the coordination stereochemistry with known Ca^2+^-specific parameters (25, 26), a marked increase in electron density relative to Na^+^, and temperature B factors consistent with those of the protein and water donor atoms (Table S2). In proteins, Ca^2+^ is generally ligated by seven oxygen atoms (neutral, carboxylate or water oxygen) in a conformation best represented by a pentagonal bipyramid (26), and this is observed for the Vn HX domain.

As predicted by NMR (24), the m1-bound Ca^2+^ is not deeply buried within the channel, and locates 0.4-0.8 Å further away from Cl^−^ than m1-bound Na^+^ (Fig. 1C-F; Fig. S1). Moreover, Ca^2+^ association results in conformational changes deep inside the channel as the m2-occluded Na^+^ shifts down further away from Cl^−^ by 0.1-0.3 Å.

The Ca^2+^ and Na^+^ structures differ at the propeller top, where the Q345 and D255 sidechains form a salt bridge in the presence of Ca^2+^ but splay apart with Na^+^ (Fig. 1A). This is accompanied by a rearrangement of the backbone at the top of the HX4 blade, and is consistent with the Ca^2+^-induced perturbations that we observed in the NMR signals of D255 and Q345 (24). The structures also differ at the bottom of the propeller, where the m2-occluded Na^+^ is stabilized either by the D166 carboxylate side chain or a sulfate anion. In the sulfate-free molecule of Vn-Ca_2_, the D166 side chain reaches deep inside the channel establishing ∼0.5 Å closer contacts between its carboxylate anion and the occluded Na^+^, resulting in a D166-capped conformation of the channel. In sulfate-bound Vn-Ca_2_, the sulfate anion arranges to form close contact with Na^+^ as well as favorable polar contacts with the backbone amide hydrogens of D166, T211, A259 and A351. This releases the K168 sidechain from the channel opening, rendering the channel bottom more accessible to solvent. The structures illustrate the extent of conformational plasticity of the HX domain and its response to Ca^2+^.

### Effect of Ca^2+^ on the Vn HX channel

Previously, we showed that Ca^2+^ binds Vn HX with a dissociation constant in the range of 27 μM (24) and, since this value is approximately ten times higher than the blood concentration of Vn (2.5-5 μM) (1-3) and thirty times lower than the total blood concentration of Ca^2+^ (1.3-1.5 mM) (26), we concluded that Vn is Ca^2+^-associated *in vivo*. Using NMR, we showed that Ca^2+^ enhances conformational exchange of Vn on the ms time scale, and by monitoring protein unfolding with DSF and an environment-sensitive fluorescent dye, we showed that Ca^2+^ markedly increases the unfolding transition temperature of Vn and, thus, protects the structure from thermal denaturation. These observations are specific for Ca^2+^ and absent for other metal ions (Na^+^, K^+^, Zn^2+^, Mg^2+^, and Ni^2+^).

To understand the molecular basis for these Ca^2+^-specific effects, we performed unrestrained all-atom MD simulations of HX, with Ca^2+^ (Vn-Ca) or Na^+^ (Vn-Na), at 30 or 60°C (Table S3). The starting structure of Vn-Ca was generated by modeling short regions of missing electron density in the Vn-Na_1_ crystal structure (PDB 6o5e, molecule A), and explicitly replacing Na^+^ with Ca^2+^ in the m1 binding site, while keeping Na^+^ at the m2 site. A 3.2 μs MD simulation shows that Ca^2+^ remains bound to the protein over this time course, with little fluctuation in its starting position at an average distance of 5.4 Å from the channel-occluded Cl^−^ anion (Fig. S2A). By contrast, an alternative model where Na^+^ was replaced with Ca^2+^ at the m2 site results in unstable association of either Ca^2+^ or Na^+^ with the m1 site (Fig. S2B). These results are in agreement with the crystal structures and previous NMR data (24) which identified m1 as the Ca^2+^-binding site.

Five independent 1 μs MD simulations of Vn-Ca or Vn-Na, result in average conformations and positions of the channel-bound ions that mirror the experimental structures (Fig. 2A-C). Ca^2+^ binds higher in the channel (∼5.4 Å from Cl^−^) than Na^+^ (4.9 Å from Cl^−^) and exhibits a narrower range of positional fluctuation at both temperatures. Ca^2+^ is also more solvent exposed and coordinates an average of 4 (30°C) or 3 (60°C) water molecules (Fig. 2D-F; Fig. S3A), compared to 1.5 water molecules associated with Na^+^ at either temperature.

**Fig. 2.**
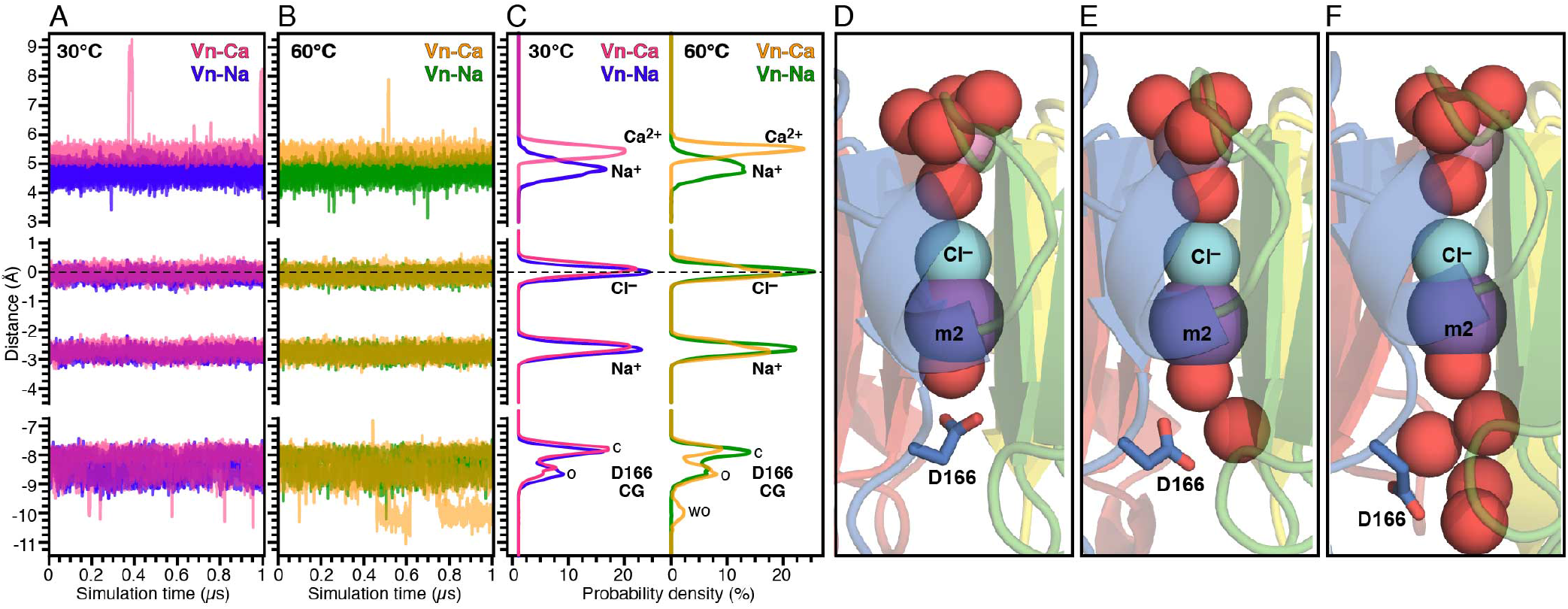
MD simulations of ion positions within the Vn HX channel. The distances of Na^+^ or Ca^2+^, Na^+^, Cl^−^ and the D166 CG atom are relative to the average position of Cl^−^ in the channel. **(A, B)** Time series of atomic positions for five independent 1 μs MD simulations. **(C)** Average probability distributions of the atomic distances extracted from the time-dependent data. Each distribution is the average over the last 500 ns of five independent 1 μs simulations. Colors denote MD data for: Vn-Ca at 30°C (pink) or 60°C (yellow), and Vn-Na at 30°C (indigo) or 60°C (green). D166 closed (c), open (o) and wide open (wo) conformations are marked. **(D-F)** MD simulation snapshots of Vn-Ca at 30°C, showing closed (D), open (E) and wide open (F) conformations of D166. Colors denote structural repeat units HX1 (blue), HX2 (green), HX3 (yellow), and HX4 (red). Sphere colors denote Na^+^ (indigo), Ca^2+^ (pink), Cl^-^ (cyan), and O (red).

At 30°C, the D166 side chain exhibits predominantly two conformations in either Vn-Ca or Vn-Na: A closed conformation (Fig. 2D) with the D166 carboxylate group capping the channel bottom as in the sulfate-free crystal structures, and an open conformation (Fig. 2E) with the channel more accessible to solvent. Increasing the temperature to 60°C equalizes the closed and open populations of D166 in Vn-Ca, and populates a third, wide-open conformation (Fig. 2F) where D166 points away from the channel. The conformational profile of Vn-Na, by contrast, is not appreciably affected by temperature.

### Ca^2+^ modulates the conformational adaptation to temperature of HX

The mean square displacement (MSD) of atomic coordinates from the starting model reflects protein flexibility, while the radius of gyration and solvent accessible surface area (SASA) provide estimates of protein compactness, all properties that are associated with protein adaptation to temperature (27, 28). The average MSD values from the four sets of MD simulations, with Ca^2+^ or Na^+^, demonstrate a distinct effect of Ca^2+^: While Vn-Na has equivalent MSD at 30°C and 60°C, the MSD of Vn-Ca is distinctly lower at 30°C but shifts within the range of Vn-Na at 60°C (Fig. 3A; Fig. S3B).

**Fig. 3.**
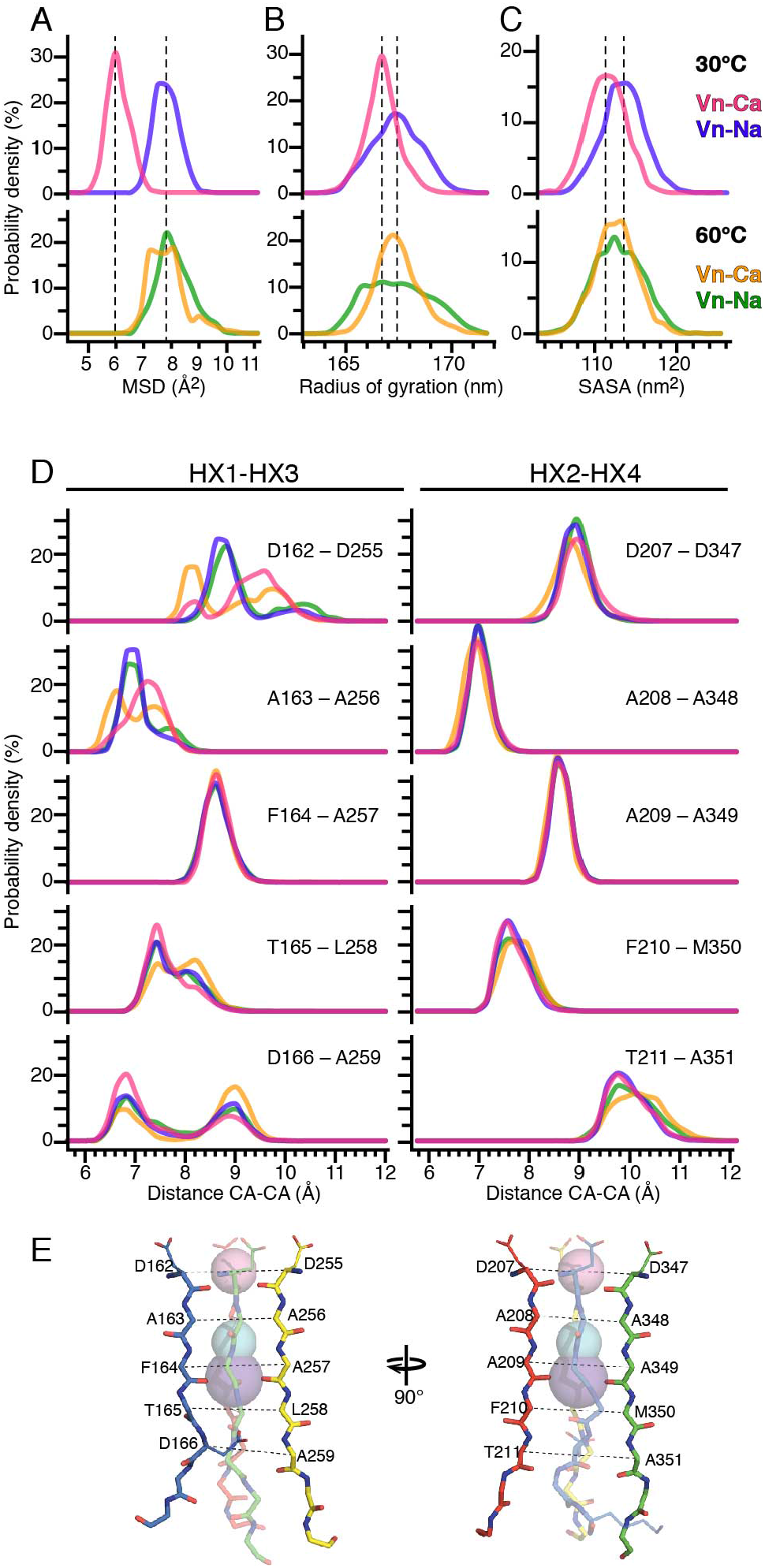
MD simulations of protein flexibility, compactness and HX channel conformation. **(A-D)** Probability distributions of MSD (A), radius of gyration (B), SASA (C), and cross-channel CA atom distances (D). Each distribution reflects the average over the last 500 ns of five independent 1 μs MD simulations. Colors denote Vn-Ca at 30°C (pink) or 60°C (yellow), and Vn-Na at 30°C (indigo) or 60°C (green). **(E)** Orthogonal views of the channel backbone in the structure of Vn-Ca_2_ (molecule A of the asymmetric unit). Colors denote structural repeat units HX1 (blue), HX2 (green), HX3 (yellow), and HX4 (red). Spheres denote Na^+^ (indigo), Ca^2+^ (pink), and Cl^-^ (cyan) ions in the channel. Dashed lines connect cross-channel HX1-HX3 and HX2-HX4 CA atoms.

Temperature also has a marked effect on the compactness of Vn-Ca, but little effect on Vn-Na (Fig. 3B, C; Fig. S3C, D): At 30°C, Ca^2+^ reduces both radius of gyration and SASA, while both parameters return to the levels of Vn-Na at 60°C, and are visibly sharper for Vn-Ca at both temperatures. The Ca^2+^-dependent temperature profile is also mirrored by the average number of water molecules found within 4 Å of the m1 metal ion: While temperature has a large effect on the water coordination of m1-bound Ca^2+^, little difference is observed in the case of Na^+^ (Fig. S3A). Overall the data indicate that Ca^2+^ enhances the protein’s adaptability to temperature, rendering it able to alter its flexibility compactness and hydration with temperature alteration.

The Ca^2+^-dependent temperature adaptability property is further reflected in the conformational response of HX to temperature. At 30°C, Ca^2+^ favors cross sectional HX1-HX3 inter-atomic distances (Fig. 3D, E) that are ∼1 Å greater than Vn-Na at the channel top, and smaller at the bottom. At the top of the channel, Ca^2+^ also broadens the inter-atomic distance distribution profile, indicating that it increases the range of conformations that the channel can sample on the μs time scale of the simulation. At 60°C, the effect of Ca^2+^ is reversed, favoring cross-sectional distances that are smaller at the channel top and greater at the bottom, and while the HX2-HX4 distances are much less susceptible to Ca^2+^ they too change slightly in the same temperature-dependent directions. By contrast, temperature has little effect on the conformation of Vn-Na, where the channel size does not vary between 30°C and 60°C.

To further examine protein conformation we calculated the number of polar contacts – including H bonds and salt bridges – mediated by no more than one water molecule. At 30°C, Ca^2+^ increases the average number of polar contacts by ∼3 compared to Na^+^, affecting predominantly residues in the channel-juxtaposed HX1 and HX3 blades (Fig. 4A, B). Examination of the MD structural models reveals that Ca^2+^ promotes the formation of contacts (D162-R176, D184-R189, E185-R353) situated longitudinally from propeller top to bottom in blades HX1 and HX3 (Fig. 4C).

**Fig. 4.**
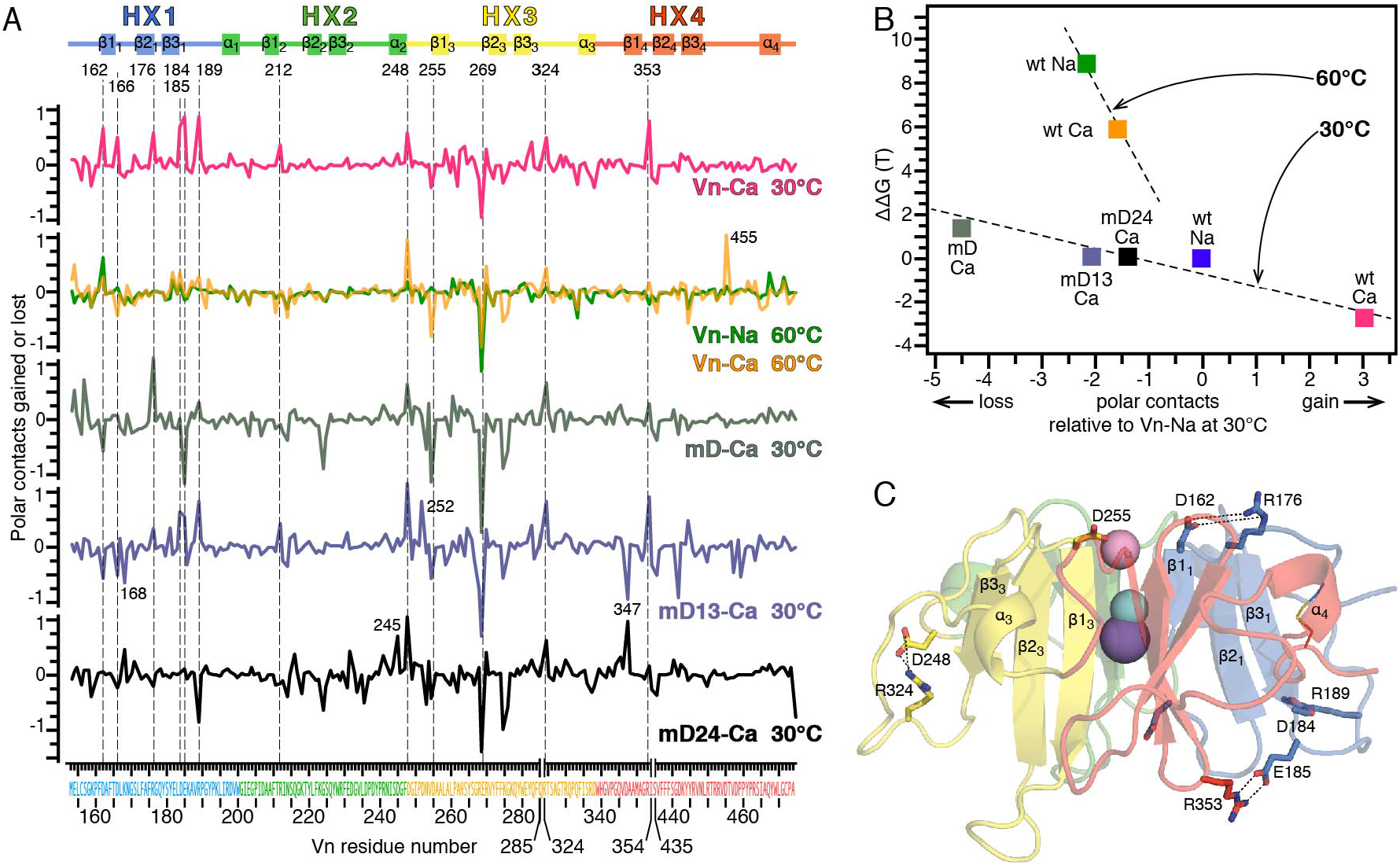
Effect of Ca^2+^ on the number of polar contacts of Vn HX from MD simulations. **(A)** Difference in number of polar contacts relative to Vn-Na at 30°C. Each trace is the average over the last 500 ns of five independent 1 μs MD simulations. Structural elements of the HX1-HX4 blades in the domain are depicted above the data. Dashed vertical lines mark residues with changes ≥0.5. (**B)** Correlation of polar contacts with values of ΔΔG estimated with eq. 1 from the experimental values of T_m_ (Table 1). Each data point is the average change in polar contacts relative to to Vn-Na at 30°C. **(C)** Key residues (sticks) with polar contacts that are affected by Ca^2+^ Colors reflect Ca^2+^-bound Vn (pink) and Na^+^-bound Vn (indigo) at 30°C. Colors denote structural repeat units HX1 (blue), HX2 (green), HX3 (yellow), and HX4 (red). Spheres denote Na^+^ (indigo), Ca^2+^ (pink), and Cl^-^ (cyan) ions in the channel.

At the bottom of the channel, Ca^2+^ favors the formation of polar contacts between the D166 carboxylate and the backbone of surrounding residues (T211, A259, A351), and this has the effect of populating the closed conformation of D166. On the other hand, Ca^2+^ disfavors contacts to the E269 sidechain, which forms salt bridges with the backbone sites of R212, I213 and F284 in the Na^+^-bound protein. Ca^2+^ also disfavors polar contact of the D255 carboxylate group to the backbone of G276. This result is in line with formation of the D255-Q345 salt bridge that is observed in the Ca^2+^-bound crystal structures.

**Table 1.** Unfolding transition temperature (T_m_) of wild-type and mutant Vn HX. Values of T_m_ reflect the average and standard deviation of at least three independent DSF experiments.

| Cation | $T_m$ (°C) | | | |
| --- | --- | --- | --- | --- |
|  | wt-Vn | mD | mD13 | mD24 |
| 0 mM $\text{Ca}^{2+}$ | 59.85 ± 0.67 | 55.71 ± 0.34 | 60.60 ± 0.60 | 58.53 ± 0.34 |
| 2 mM $\text{Ca}^{2+}$ | 66.82 ± 0.64 | 56.11 ± 0.34 | 60.55 ± 0.51 | 59.71 ± 0.68 |

Finally, Ca^2+^ association induces the formation of a water-bridged network, involving the rim-Asp carboxylates and the Ca^2+^ cation, in agreement with the crystal structures. This water network exchanges less rapidly with bulk solvent on the μs timescale of MD simulation. Increasing the temperature to 60°C reduces the number of polar contacts overall, but Ca^2+^ continues to offer advantage over Na^+^.

The enrichment of polar interactions is a key source of higher folding stability and lower heat capacity of unfolding in thermophilic proteins (29-31). In the case of Vn HX, both crystal structures and MD simulations indicate that Ca^2+^ increases the average number of direct and water-mediated polar contacts relative to Na^+^. To assess the effect on the protein’s thermodynamic stability we estimated the difference in unfolding free energy (ΔΔG) between the Ca^2+^ and Na^+^ bound states, using the generic stability curve equation (Eq. 1) developed by Rees and Robertson (31): 

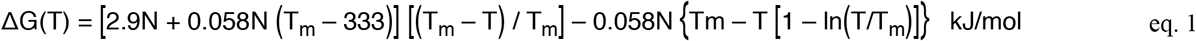

where ΔG(T) is the free energy of unfolding at a given temperature T, N is the number of protein residues and T_m_ is the temperature of unfolding. Based on the experimental values of T_m_ that we measured for Vn HX (204 residues) with Ca^2+^ or Na^+^ (Table 1), the estimates of ΔG(T) indicate that Ca^2+^ confers 2.7 kcal/mol of extra stability at 30°C, and 3.0 kcal/mol of stability at 60°C, over Na^+^ (Fig. 4B). These estimates are within the range determined for thermophilic proteins that acquire thermal stability by increasing their polar contacts (31).

### Role of rim-Asp in Ca^2+^-induced stabilization

To understand which residues of Vn are important for Ca^2+^-specific stabilization, we compared its amino acid sequence (Fig. S4) with those of the HX domains of two matrix metalloproteases (mmp) – mmp-14 which does not bind Ca^2+^ (32) and mmp-19 which binds Ca^2+^ and also gains thermal stability (33). In the sequence of mmp-14, the rim-Asp of repeats HX2 and HX4 are substituted with N and G, while those of mmp-19 are conserved with the exception of HX4 where D is replaced by S. Moreover, mmp-19 shares other amino acid identities with Vn residues that form polar contacts in the Ca^2+^-bound protein (R176, E185, D248, R324 and Q345).

Based on this analysis, we performed MD simulations of three Vn HX mutants: One where all four rim-Asp were mutated to Asn (mutant mD), and two where the rim-Asp were mutated as channel-juxtaposed pairs in HX1 and HX3 (mutant mD13: D162N and D255N) or HX2 and HX4 (mutant mD24: D207N and D347N). Five independent MD simulations for each mutant, at 30°C, indicate that Ca^2+^ remains associated with the protein over the course of 1 μs (Fig. S5). In all mutants, Ca^2+^ binds slightly higher above the channel (∼0.1 Å) than wild-type, and the occluded ions and D166 sidechain exhibit a broader range of conformational dynamics in the mD24 and mD mutants. These mutants also exhibit broader ranges of flexibility and compactness parameters than wild-type Vn-Ca (Fig. S3). Notably, the number of polar contacts in all three mutants is either slightly lower (mD13, mD24) or much lower (mD) than wild-type Vn-Na, and appreciably lower than wild-type Vn-Ca (Fig. 4A, B). Taken together, the MD data suggest that the rim-Asp are important for the Ca^2+^-induced stabilization of Vn.

To assess the effects of these mutations experimentally, we purified them and examined their conformational fitness, Ca^2+^ binding ability and thermodynamic stability by means of NMR and DSF. The NMR spectra show that the mutations do not disrupt the global fold of the protein, notwithstanding minor differences from residues near the mutation sites (Fig. 5A-D; Fig. S5). The spectra of mD13 and mD24 have narrow, homogeneous lines and are well-resolved. The addition of Ca^2+^ induces chemical shift perturbations similar to those observed for wild-type, and thus consistent with Ca^2+^ association at the top of the channel, indicating that mD13 and mD24 have not lost their Ca^2+^ binding ability, in line with the MD simulations. By contrast, the NMR spectrum of mD has very broad, poorly resolved lines, that reflect a high degree of conformational exchange on the ms time scale. Its general resemblance to the wild-type spectrum suggests that the overall fold is maintained, but its high propensity for precipitation precluded Ca^2+^ binding analysis by NMR.

**Fig. 5.**
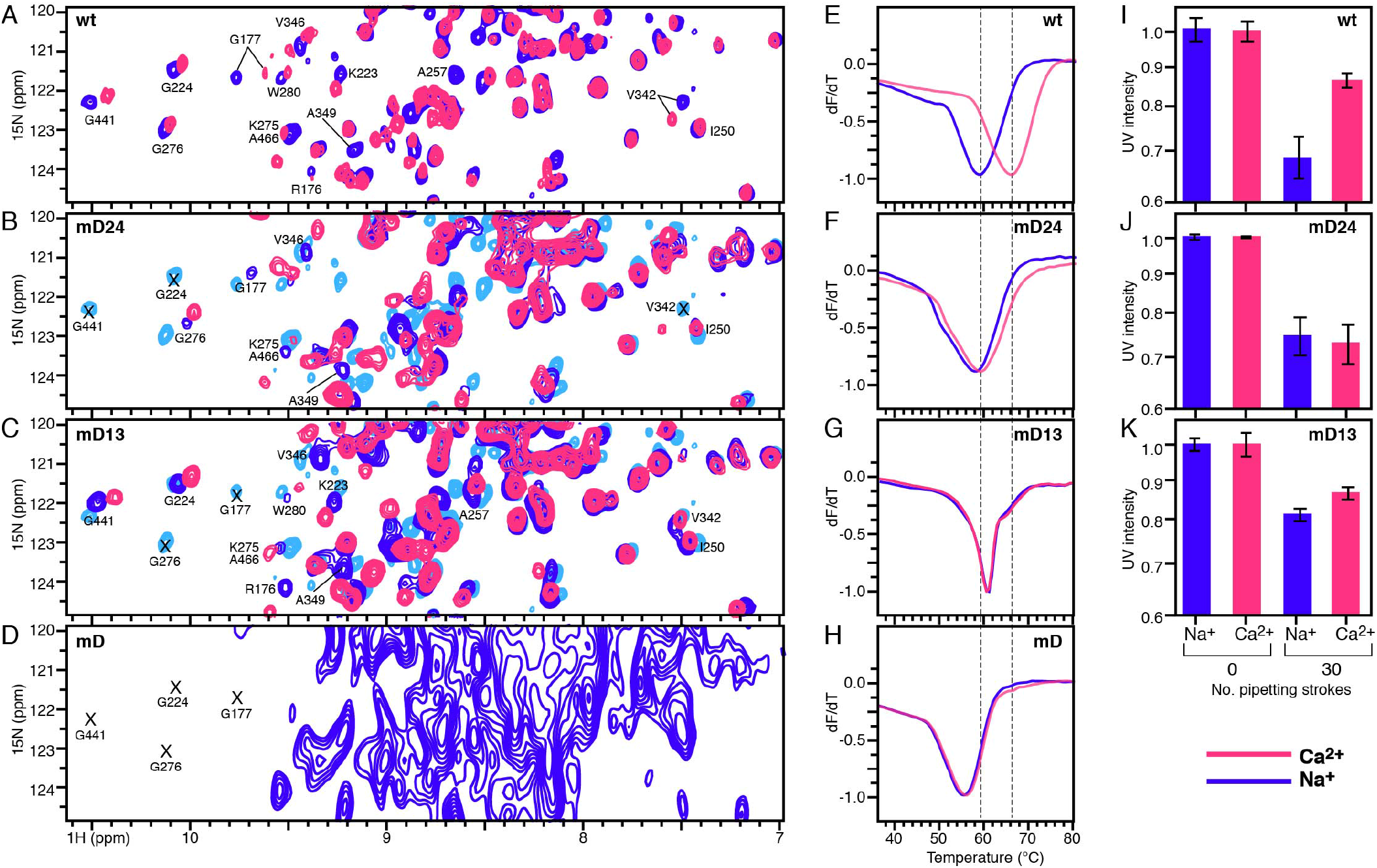
Effect of rim-Asp mutations. NMR, DSF and shear stress analysis data for wt and mutant Vn HX, without (blue) or with (pink) 2 mM CaCl_2_. **(A-D)** Selected regions of 2D solution NMR ^1^H/^15^N HSQC spectra. Missing peaks from Gly residues are marked (X) **(E-H)** Representative DSF traces. **(I-K)** Shear stress analysis. The intensity of UV absorbance at 280 nm reflects the % protein remaining in solution after 60 passes through the 0.21 mm diameter needle. Error bars reflect standard deviation for 6-18 independent experiments.

Interestingly, DSF analyses (Fig. 5E-H, Table 1) show that while wild-type is substantially stabilized by 2 mM Ca^2+^, the three mutants are essentially not susceptible to Ca^2+^, and in the case of mD, have lower (mD) thermal stability than wild-type in Na^+^. All three mutants, therefore, appear to have lost the Ca^2+^-induced thermal adaptability property, even though mD13 and mD24 retain the ability to bind Ca^2+^. These results correlate with the loss of polar contacts predicted by the MD simulations for the mutants (Fig. 4A, B).

### Ca^2+^ protects Vn from shear stress

Finally, we sought to explore the potential significance of the Ca^2+^ association for Vn in physiology, where extreme temperature excursions like those tested in this study do not occur, and temperature is maintained within the narrow homeostatic range near 37°C. In the vasculature, proteins are subject to hydrodynamic shear rates of 500-1,000 sec^-1^ (34) and hydrodynamic flow can induce protein remodeling, unfolding and aggregation. In this setting, the ability to adapt to environmental changes in shear force acquires importance. For example, von Willebrand factor (VFW), which also circulates in the high-shear environment of the vasculature, binds Ca^2+^ via its A2 domain to resist shear-induced unfolding and cleavage by the metalloprotease ADAMTS-13 (35, 36).

To test whether Ca^2+^ can protect Vn HX against shear stress, we performed an assay where we estimated the amount of protein that aggregates out of solution after multiple passes through a narrow diameter (0.21 mm) needle, as described (37). The 12 μM (0.25 μg/ml) protein solution is in the range of the physiological concentration of Vn (0.2–0.5 mg/ml), and we estimate that each pass of 150 μL s^-1^ through the needle, generates a shear rate of ∼165,000 s^-1^ (38), much greater than the hydrodynamic shear rates in human vasculature, although the experimental time exposure to shear force is much lower than the continuous hydrodynamic flow in the vasculature. After 60 passes through the needle (Fig. 5I-K), the soluble concentration of wild-type HX, estimated by measuring UV absorbance at 280 nm, is ∼20% greater with Ca^2+^ than Na^+^, while no changes were observed when the protein solutions were incubated without pipetting in the spectrophotometer cuvette, over the same time period. In the mD24 and mD13 mutants, by contrast, the level of soluble protein does not appear to be affected by Ca^2+^, and is reduced to an approximately equal extent in either Ca^2+^ or Na^+^ after shearing. The data mirror the response to temperature: Ca^2+^ cooperates with the four rim-Asp to protect Vn HX against shear stress.

## Conclusions

In summary, we have resolved the Ca^2+^-bound structure of the Vn HX domain and described how Ca^2+^ enables Vn HX to adapt to large excursions of temperature and mechanical shear force. The crystal structures and MD simulations indicate that Ca^2+^ stabilizes multiple conformational states, enabling the protein to alter its conformation as dictated by environmental conditions. The four rim-Asp that form an electronegative corral around the Ca^2+^ binding site are not absolutely required for Ca^2+^ binding, but they play an important in the Ca^2+^-induced adaptability to temperature and shear stress through their participation in a network of water-mediated and direct polar contacts with Ca^2+^ and other sidechains. Ca^2+^ promotes a reorganization of polar contacts that relays from top to bottom of the HX propeller and is accompanied by a conformational change of the channel. Our results indicate that the hydration sphere of Ca^2+^ plays an important role in the environmental adaptation of Vn HX.

The data suggest a model for the way in which Vn HX adapts to its environment. At 30°C (Fig. 6A), the top of the β1 strands splay apart to accommodate a Ca^2+^ ion and the additional water molecules in its coordination sphere. This results in a conformational rearrangement of polar contacts that propagates from the Ca^2+^ binding site at the top of the channel, around the propeller exterior, and all the way to the channel bottom. At 60°C by contrast (Fig. 6B), increased exchange of the Ca^2+^ hydration shell with bulk water reduces the effective number of water molecules that coordinate Ca^2+^. The loss of water in the Ca^2+^ coordination sphere is compensated by the electronegative rim-Asp carboxylates, which rearrange to make closer contact with the positively charged Ca^2+^ center, causing the top of the channel to constrict. In the rim-Asp mutants this compensatory effect is impaired, and they lose the ability to adapt to temperature and mechanical stress. While the precise types of conformational changes that operate under shear remais to be determined, the Ca^2+^-induced adaptability property is likely to play an important role for the activity of Vn in blood.

**Fig. 6.**
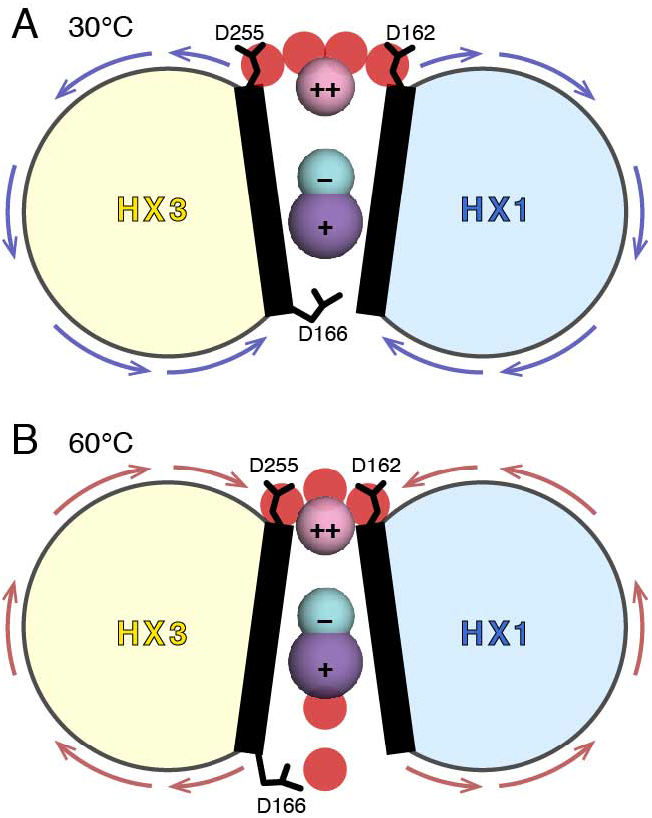
Model for the Ca2+-mediated temperature adaption of Vn HX. **(A)** Adaptability to 30°C. **(B)** Adaptability to 60°C.

Unlike the large rearrangements observed for archetypal Ca^2+^ binding proteins, such as concanavalin, calmodulin and troponin C (26), the Ca^2+^-induced conformational changes observed for the HX domain of Vn are subtle. Nevertheless, the formation of just a few polar contacts arising from small rearrangements has been shown to result in sufficient differences in stabilization free energy that protect proteins against extremes of temperature and pressure (29-31). For example, a stabilization free energy difference in the range of 5 kcal/mol can be readily attained by forming a few polar H bonds or as little as two salt bridges.

Ca^2+^ appears to be the ideally suited for promoting a protein’s adaptability to environment. Its valency, polarizability and ionic radius, as well as its hydration energy, hydrated radius and related charge density, all make Ca^2+^ highly adaptable to sites of irregular coordination geometry, such as those of the Vn HX domain (26). The coordination flexibility of Ca^2+^ is reflected in its variable coordination number (typically 6-8 and up to 12), coordination geometry and bond lengths (2.2-2.4 Å). By contrast, the coordination geometry of Na^+^ is less flexible because its lower charge density (∼2 times less than Ca^2+^) and hydration energy (∼4 times less than Ca^2+^) restrict the coordination number (typically 4-6 with octahedral geometry).

Vn is a known regulator of osteoclast bone resorption (39), and Ca^2+^ has been shown to play a role in the interaction of Vn with the leukocyte integrin α_M_β_2_ (Mac-1), independent of either the Vn RGD motif or somatomedin-B domain (40). Moreover, the association of Vn with Ca^2+^ is central in the ectopic calcified protein-lipid deposits associated with age-related macular degeneration and other degenerative pathologies. We propose that Ca^2+^ plays an important role in adapting Vn to the diverse environments where it functions.

## Materials and Methods

### Protein preparation

The HX domain sequence includes Vn residues 154-285, 324-354 and 435-474. The methods for *E. coli* protein expression and preparation were as described (23). Mutations were introduced by

### Crystallization, X-ray data acquisition and structure determination

A solution of HX (6 mg/mL) was dialyzed overnight into buffer A (20 mM MES pH 6.5, 100 mM NaCl, and 1 mM CaCl_2_), and then cleared by centrifugation. The final protein concentration was 4 mg/mL. Crystals were obtained by mixing 0.2-0.3 μL of Vn HX (4 mg/mL in buffer A) with 0.15 μL of precipitant solution [20 mM MES pH 6.5, 90 mM imidazole, 30 mM NaNO_3_, 30 mM Na_2_HPO_4_, 30 mM (NH_4_)_2_SO_4_, 11.25% (v/v) 2-methyl-2,4-pentanediol, 11.25% (w/v) PEG-1000, 11.25% (w/v) PEG-3350, and 3% (w/v) d-(+)-trehalose], and equilibration in a sitting drop plate with 50 μL of precipitant solution at room temperature. Crystals appeared within a week and grew for additional 2-3 weeks. To analyze metal ion binding, the crystals were soaked overnight with 1, 100 or 200 mM CaCl_2_, or 50 mM KCl in Na^+^-free soaking solution [20 mM Bis-Tris methane-Cl pH 6.5, 70 mM NH_4_NO_3_, 11.25% (v/v) 2-methyl-2,4-pentanediol, 11.25% (w/v) PEG-1000, 11.25% (w/v) PEG-3350, and 3% (w/v) d-(+)-trehalose]. Soaking with KCl solution destroyed the crystals. The CaCl_2_-soaked crystals were flesh frozen in liquid nitrogen and shipped to the Stanford Synchrotron Radiation Lightsource (SSRL) for data collection.

X-ray diffraction data were collected at the SSRL beamline BL12-2, at a wavelength of 0.97946 Å and temperature of 100 K. The data were processed using the CCP4 suite (41). The structure of Vn-Ca_1_ was solved first by molecular replacement of the previously published Vn-Na_1_ structure (PDB code 6o5e; 100% identity), and then used to solve other structures. Phenix.AutoBuild (42) was used for initial model building, followed by several rounds of manual model inspection and correction in Coot (43) and refinement by phenix.refine (42) and Refmac5 (44). Comparison of the structures soaked with various CaCl_2_ concentrations did not reveal significant differences in binding of metal ions. The data collection and refinement statistics are presented in the Table 1. The structure coordinates were deposited in the protein data bank (PDB) with accession codes 7txr (Vn-Ca_1_), 7rj9 (Vn-Ca_2_) and 7u68 (Vn-Na_2_).

Molprobity (45) and the PDB validation server were used for structure validation throughout refinement. All structures had Ramachandran statistics with more than 95% of residues in favored positions and less then 1% outliers.

### NMR spectroscopy

The NMR samples were prepared as described (23). NMR experiments were performed at 30°C, on a Bruker Avance 600 MHz spectrometer equipped with a ^1^H/^13^C/^15^N triple-resonance cryoprobe. The NMR data were processed and analyzed using TopSpin (Bruker). Assignments of the ^1^HN and ^15^N chemical shifts were transferred from the previously assigned data for HX with Na^+^ or Ca^2+^ (23) (BMRB codes 50241 and 50261).

### DSF experiments

Protein melting experiments were performed in a 96-well plate format, using a LightCycler 480 instrument (Roche), with a linear temperature gradient of 0.03°C/sec from 20°C to 95°C as described previously (23).

### Shear stress experiments

Methods were adapted from to published protocols (37, 46). Protein solutions were prepared in a 150 μL volume, with 12 μM Vn HX, 20 mM MES (pH 6.5), 30 mM NaCl, and an additional 8 mM NaCl or 8 mM CaCl_2_. Shear force was applied using a 27 G needle with internal diameter of 0.21 mm, and length of 12.7 mm, fitted to a 1 mL syringe. In each 1 sec syringe stroke, the protein solution was drawn and extruded through the needle to produce ∼750 dynes/cm^2^ per stroke. After 30 repetitive syringe strokes, insoluble protein aggregates were removed by centrifugation (17,000 g, 5 min) and analyzed by SDS-PAGE, and the remaining soluble protein concentration was estimated by measuring UV absorbance (280 nm).

### MD Simulations

All-atom MD simulations were performed as described (47), using the CHARMM36 force fields (48) with the TIP3P water model (49). The temperature was set to 303.15K or 333.15K for simulations at 30°C or 60°C, and the pressure was maintained at 1 bar.

All systems for MD (Table S3) were prepared and equilibrated using CHARMM-GUI *Solution Builder* (50). The initial structural models were generated from the crystal structure of Vn-Na_1_ (PDB code 6o5e, molecule A). All ligands and water molecules were removed, and residues with missing electron density were modeled using GalaxyFill (51). For the Ca^2+^-bound simulations, Na^+^ ions were replaced with Ca^2+^, the models were solvated in a cubic water box, and then equilibrated with the CHARMM-GUI standard protocol for 125 ps, before the production runs.

MD production simulations were conducted with OpenMM (52) for 1 μs, and the last 500 ns of trajectories were used for analysis. Five 1 μs MD simulations were performed for each of four wild-type Vn HX bound to Na^+^ or Ca^2+^ (Vn-Na and Vn-Ca) at 30°C or 60°C, and for each of three mutants bound to Ca^2+^ (mD-Ca, mD13-Ca and mD24-Ca) at 30°C, for a total of 35 independent simulations (Table S3).

Four additional 3.2 μs simulations were performed for each of four wild-type Vn HX systems with different Na^+^ and Ca^2+^ ion combinations at channel positions m1 and m2, at 30°C.

For each simulation, trajectories were generated every 1 ns, and trajectories from all five simulations were combined for analysis in each system. Statistical analyses were performed with home-made Python scripts in MDAnalysis and JupyterNotebook.

## Supporting Information

This article contains supporting information online

## Author Contributions

YT, KS, AEA, WI and FMM performed research

YT, KS, AEA, WI and FMM analyzed the data

YT and FMM wrote the paper

YT, WI and FMM designed the research

## Data and Materials Availability

All data needed to evaluate the conclusions are present in the paper and/or the Supplementary Materials. Additional data are available upon request.

## Acknowledgements

This study was supported by grants from the National Institutes of Health (GM 118186, CA030199), the National Science Foundation (MCB1810695), and the Canadian Institutes of Health Research (201711MFE-395794-210656)

