## Supplementary Tables and Figures for "Calcium-induced environmental adaptability of the blood protein vitronectin"

**Table S1. Data collection and refinement statistics (molecular replacement).** [a, b]

| | **Vn-Ca$_1$ (PDB code 7txr)** | **Vn-Ca$_2$ (PDB code 7rj9)** | **Vn-Na$_1$ (PDB code 6o5e)[c]** | **Vn-Na$_2$ (PDB code 7u68)** |
|---|---|---|---|---|
| Data collection | | | | |
| Space group | $P2_1$ | $P2_1$ | $P\ 2_1$ | $P\ 2_1$ |
| Cell dimensions | | | | |
| *a*, *b*, *c* (Å) | 40.84, 98.28, 49.43 | 0.83, 125.57, 40.88 | 40.58 124.90 40.78 | 40.56 97.36 49.16 |
| $\alpha, \beta, \gamma$ (°) | 90, 99.80, 90 | 90, 119.25, 90 | 90, 119.50, 90 | 90 99.90 90 |
| Resolution (Å) [b] | 37.26 – 2.00 (2.07 – 2.0) | 35.67 – 1.7 (1.73 – 1.7) | 35.49 – 2.00 (2.05 – 2.0) | 26.96–1.90 (1.94–1.90) |
| No. of reflections | 23890 (1561) | 38571 (1742) | 23279 (1712) | 28986 (2453) |
| Wavelength (Å) | 0.97946 | 0.97946 | 0.97946 | 1.54 |
| $R_{merge}$ [b] | 0.122 (0.523) | 0.061 (0.59) | 0.082 (0.73) | 0.071 (0.33) |
| $\langle I / \sigma(I) \rangle$ [b] | 5.6 (1.6) | 10.6 (1.6) | 9.9 (2.0) | 7.2 (2.0) |
| $CC_{1/2}$ [b] | 0.84 (0.72) | 0.99 (0.65) | 0.99 (0.73) | ND |
| Completeness (%) [b] | 92.0 (60.6) | 97.9 (84.9) | 97.6 (94.8) | 0.98 (0.96) |
| Redundancy [b] | 3.3 (2.6) | 3.5 (2.8) | 3.5 (3.3) | 2.3 (2.3) |
| Refinement | | | | |
| Resolution (Å) | 37.26 – 2.00 | 35.67 – 1.7 | 35.52 – 2.00 | 24.21 – 1.90 |
| No. reflections / test set | 23887 (1560) | 38521 (2258) | 21837 (1419) | 28985 (1472) |
| $R_{work}$ / $R_{free}$ | 0.184 / 0.226 | 0.17 / 0.21 | 0.184 / 0.247 | 0.195 / 0.230 |
| No. atoms | | | | |
| Overall | 3207 | 3567 | 3364 | 3282 |
| Protein | 3019 | 3212 | 3162 | 3079 |
| Ligand / Ion | 25 | 11 | 31 | 30 |
| Water | 163 | 344 | 171 | 173 |
| Average B-factors | | | | |
| Overall | 44.70 | 25.97 | 34.77 | 35.32 |
| Protein | 44.66 | 24.96 | 34.57 | 35.32 |
| Ligand / Ion | 51.32 | 19.62 | 31.97 | 41.21 |
| Water | 44.92 | 35.67 | 39.09 | 36.56 |
| R.M.S. deviations from ideal | 0.013 | 0.013 | 0.013 | 0.009 |
| Bond lengths (Å) | 1.69 | 1.74 | 1.70 | 1.43 |
| Bond angles (deg) | | | | |
| Ramachandran plot | 97.43 | 97.57 | 97.82 | 99.16 |
| Favored (%) | 0.29 | 0.00 | 0.00 | 0.0 |
| Outliers (%) | | | | |

[a] Each data set was collected from a single crystal.

[b] Values in parentheses are for the highest-resolution shell.

[c] Previously published PDB entry 6o5e was re-refined with RefMac.

**Table S2. Atomic parameters of the channel structures.** Inter-atomic distances (D) and B factors (B) in the coordination spheres of the $Ca^{2+}$ and $Na^{+}$ metal ions bound at the m1 and m2 channel sites.

| | **$Ca^{2+}$-bound Vn-HX ($m1=Ca^{2+}$)** | | | | | | | | **$Na^{+}$-bound Vn-HX ($m1=Na^{+}$)** | | | | | | | |
|---|---|---|---|---|---|---|---|---|---|---|---|---|---|---|---|---|
| | **Vn-$Ca_1$ (PDB code 7txr)** | | | | **Vn-$Ca_2$ (PDB code 7rj9)** | | | | **Vn-$Na_1$ (PDB code 6o5e)** | | | | **Vn-$Na_2$ (PDB code 7u68)** | | | |
| | **mol A** | | **mol B** | | **mol A** | | **mol B** | | **mol A** | | **mol B** | | **mol A** | | **mol B** | |
| **Metal coordination atom** | **D (Å)** | **B (Å$^2$)** | **D (Å)** | **B (Å$^2$)** | **D (Å)** | **B (Å$^2$)** | **D (Å)** | **B (Å$^2$)** | **D (Å)** | **B (Å$^2$)** | **D (Å)** | **B (Å$^2$)** | **D (Å)** | **B (Å$^2$)** | **D (Å)** | **B (Å$^2$)** |
| m1 | | 26.96 | | 34.36 | | 14.46 | | 10.73 | | 16.75 | | 20.67 | | 24.44 | | 21.18 |
| H2O (O) | – | | 2.8 | 37.15 | – | – | – | – | – | – | – | – | – | – | – | – |
| H2O (O) | 2.3 | 31.37 | 2.1 | 42.70 | 2.5 | 18.68 | 2.6 | 20.06 | – | – | – | – | – | – | – | – |
| H2O (O) | 2.4 | 45.10 | 2.3 | 54.96 | 2.5 | 15.62 | 2.4 | 19.17 | – | – | – | – | – | – | – | – |
| H2O (O) | 2.5 | 25.98 | 2.6 | 45.93 | 2.5 | 22.47 | 2.4 | 30.36 | 2.1 | 25.41 | 2.1 | 27.46 | – | – | – | – |
| Asp162 (O) | 2.4 | 27.17 | 2.2 | 31.75 | 2.4 | 17.02 | 2.3 | 12.52 | 2.3 | 16.51 | 2.2 | 20.04 | 2.2 | 26.00 | 2.2 | 18.35 |
| Asp207 (O) | 2.3 | 27.34 | 2.2 | 34.35 | 2.3 | 13.47 | 2.3 | 10.41 | 2.3 | 20.25 | 2.2 | 23.46 | 2.2 | 23.41 | 2.2 | 19.49 |
| Asp255 (O) | 2.4 | 24.11 | 2.3 | 36.19 | 2.4 | 15.33 | 2.3 | 12.12 | 2.3 | 20.68 | 2.2 | 21.69 | 2.1 | 21.27 | 2.0 | 17.52 |
| Asp347 (O) | 2.3 | 29.38 | 2.4 | 36.68 | 2.3 | 14.38 | 2.4 | 12.49 | 2.3 | 21.81 | 2.4 | 22.70 | 2.1 | 22.22 | 2.2 | 21.49 |
| $Cl^{-}$ | 5.4 | 23.88 | 5.5 | 34.13 | 5.5 | 13.89 | 5.5 | 12.88 | 5.1 | 16.84 | 5.2 | | 4.7 | 22.71 | 4.9 | 18.45 |
| $m2 = Na^{+}$ | | 21.98 | | 35.00 | | 11.94 | | 11.29 | | 14.20 | | 19.94 | | 20.98 | | 15.00 |
| $Cl^{-}$ | 2.8 | 23.88 | 2.8 | 34.13 | 2.8 | 13.89 | 2.9 | 12.88 | 2.7 | 16.84 | 2.8 | 20.20 | 3.1 | 22.71 | 3.0 | 18.45 |
| Phe164 (O) | 2.3 | 21.68 | 2.3 | 36.29 | 2.3 | 17.43 | 2.3 | 12.94 | 2.3 | 17.51 | 2.3 | 18.02 | 2.5 | 20.14 | 2.2 | 20.14 |
| Ala209 (O) | 2.2 | 21.06 | 2.3 | 30.57 | 2.3 | 12.46 | 2.3 | 11.05 | 2.2 | 14.82 | 2.3 | 18.78 | 2.3 | 19.75 | 2.2 | 15.14 |
| Ala257 (O) | 2.2 | 23.09 | 2.3 | 33.74 | 2.3 | 13.86 | 2.4 | 11.11 | 2.3 | 19.63 | 2.2 | 18.02 | 2.4 | 20.43 | 2.3 | 16.70 |
| Ala349 (O) | 2.3 | 22.21 | 2.3 | 31.57 | 2.3 | 15.43 | 2.3 | 12.70 | 2.3 | 14.37 | 2.2 | 19.74 | 2.4 | 20.73 | 2.4 | 18.35 |
| H2O (O) | 2.3 | 50.17 | – | – | – | – | – | – | 2.3 | 25.29 | – | – | | | | |
| $SO_4^{2-}$ (O) | – | – | 2.8 | 46.75 | – | – | 2.6 | 27.35 | – | – | 2.8 | 37.04 | 2.5 | 28.52 | 2.4 | 23.74 |
| D166 (OD1) | 4.8 | 35.39 | – | – | 4.3 | 24.53 | – | – | 4.8 | 22.39 | – | – | – | – | – | – |
| D166 (OD2) | 5.2 | 35.22 | – | – | 5.0 | 22.76 | – | – | 5.2 | 24.47 | – | – | – | – | – | – |

**Table S3. MD simulation systems.**

| Vn-HX System | Bound cation configuration | | MD time (μs) | No. of MD simulations | Temperature (°C) | System size | No. of atoms | No. of water molecules | No. of ions | | |
|---|---|---|---|---|---|---|---|---|---|---|---|
| | m1 | m2 | | | | | | | $Ca^{2+}$ | $Na^{+}$ | $Cl^{-}$ |
| Vn-Ca | $Ca^{2+}$ | $Na^{+}$ | 1 | 5 | 30 and 60 | 84 x 84 x 84 | 17,672 | 17,672 | 9 | 103 | 123 |
| Vn-Na | $Na^{+}$ | $Na^{+}$ | 1 | 5 | 30 and 60 | 84 x 84 x 84 | 17,681 | 17,681 | 8 | 104 | 122 |
| wt CaCa | $Ca^{2+}$ | $Ca^{2+}$ | 3.2 | 1 | 30 | 84 x 84 x 84 | 17,672 | 17,672 | 9 | 103 | 123 |
| wt NaCa | $Na^{+}$ | $Ca^{2+}$ | 3.2 | 1 | 30 | 84 x 84 x 84 | 17,681 | 17,681 | 8 | 104 | 122 |
| wt CaNa | $Ca^{2+}$ | $Na^{+}$ | 3.2 | 1 | 30 | 84 x 84 x 84 | 17,672 | 17,672 | 9 | 103 | 123 |
| wt NaNa | $Na^{+}$ | $Na^{+}$ | 3.2 | 1 | 30 | 84 x 84 x 84 | 17,681 | 17,681 | 8 | 104 | 122 |
| mD13-Ca | $Ca^{2+}$ | $Na^{+}$ | 1 | 5 | 30 | 70 x 70 x 70 | 9,728 | 9,728 | 10 | 57 | 81 |
| mD24-Ca | $Ca^{2+}$ | $Na^{+}$ | 1 | 5 | 30 | 98 x 98 x 98 | 28,711 | 28,711 | 12 | 167 | 195 |
| mD-Ca | $Ca^{2+}$ | $Na^{+}$ | 1 | 5 | 30 | 70 x 70 x 70 | 9,719 | 9,719 | 10 | 57 | 83 |

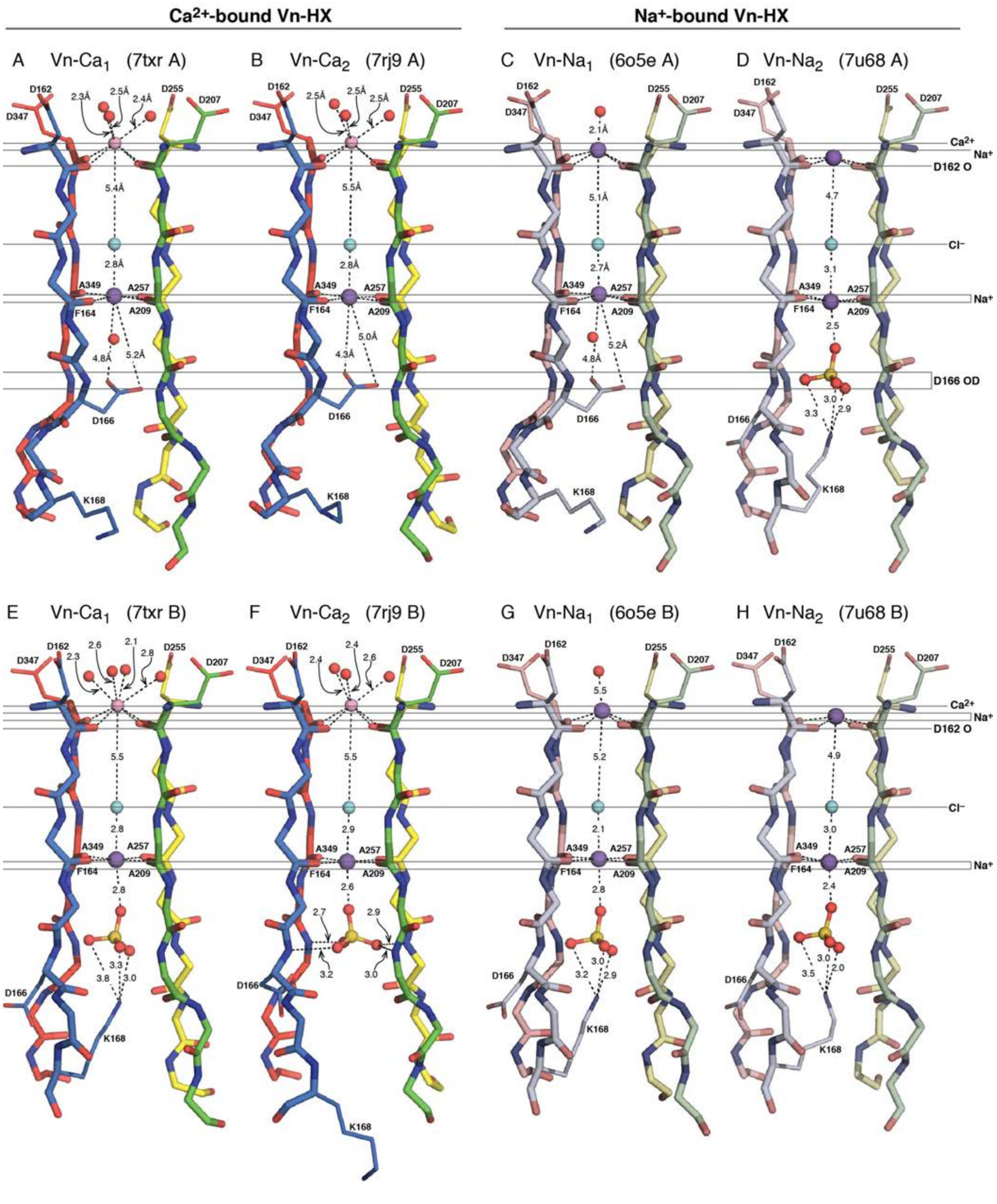

**Figure S1. Backbone structures of the Vn HX channel with $Ca^{2+}$ or $Na^{+}$ bound at the m1 site.**

**(A-D)** Molecules A and **(E-H)** molecules B of the asymmetric unit are shown for each structure. Colors denote structural repeat units HX1 (blue), HX2 (green), HX3 (yellow) and HX4 (red). Spheres denote $Na^{+}$ (indigo), $Ca^{2+}$ (pink), $Cl^{-}$ (cyan), and O (red) or S (yellow) atoms from water or bound $SO_4^{2-}$. Horizontal lines mark the positions of occluded ions and the D166 carboxylate.

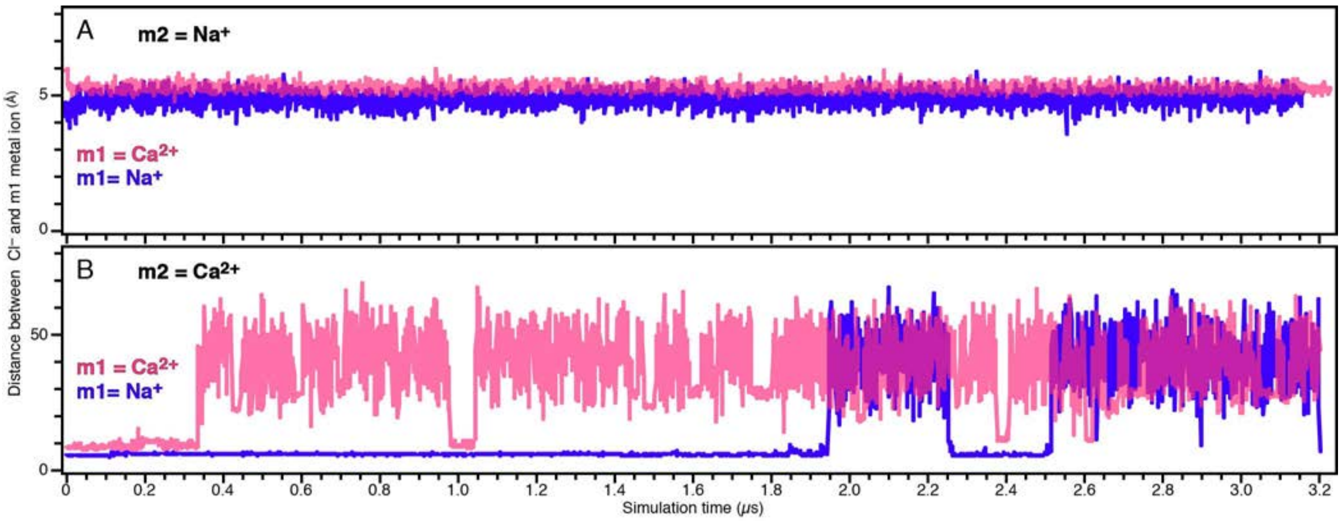

**Figure S2. Occupancy of $Ca^{2+}$ or $Na^{+}$ ions at the m1 channel site.**

**(A, B)** MD simulation time series of the distance between the channel-occluded $Cl^{-}$ anion and the $Ca^{2+}$ (pink) or $Na^{+}$ (indigo) cation initially placed in the channel m1 site, with either $Na^{+}$ (A) or $Ca^{2+}$ (B) in the m2 site. Each time series is one independent MD simulation at 30°C. Note the 10-fold scale difference between panels A and B.

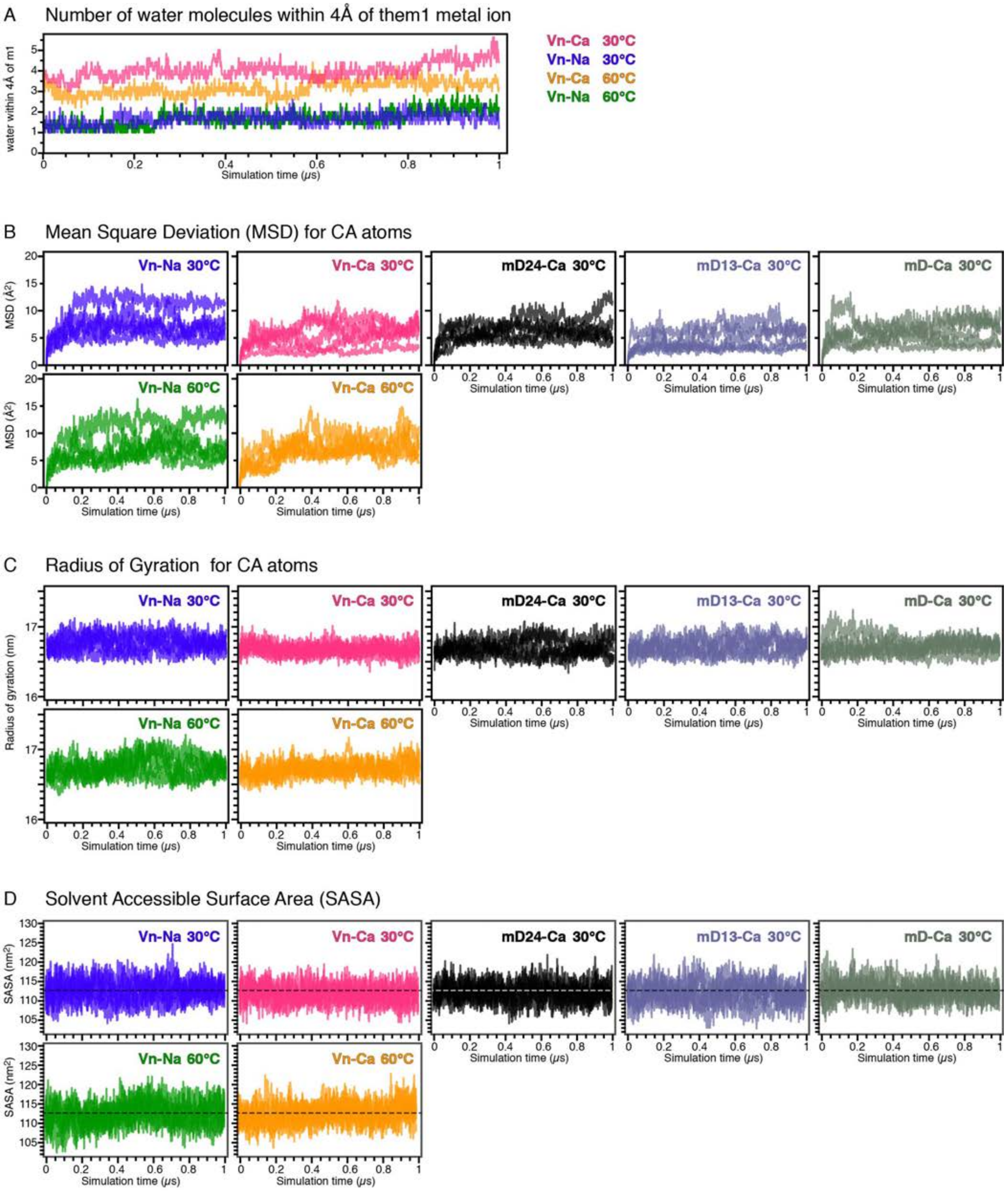

**Figure S3. MD simulations of metal ion hydration, protein flexibility and compactness.**

**(A)** Time series of number of water molecules within 4 Å of $Ca^{2+}$ or $Na^{+}$ bound at the m1 site. Each trace is the average of five independent 1 µs MD simulations.

**(B-D)** Time series of MSD (B), radius of gyration (C), and SASA (D) for wild-type Vn HX or mutants mD24, mD13 and mD. Each trace is one of five independent 1 µs MD simulations.

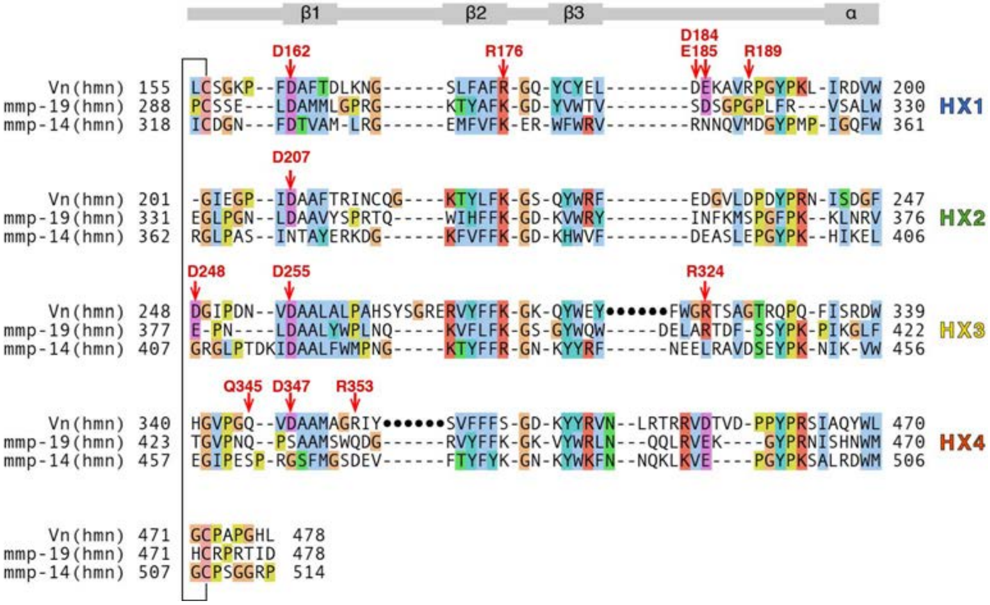

**Figure S4. Structure-based sequence alignment of the HX domains of Vn, mmp-19 and mmp-14.**

Key conserved residues are marked (red arrows). Black circles denote Vn insertion regions excluded from the alignment. The four HX repeats are marked on the right. The disulfide linked Cys residues that circularize each propeller are connected on the left. Conserved secondary structure is shown above the sequences. Alignments were generated and colored with Clustal using Jalview (Clamp et al., 2004, *Bioinformatics* 20: 426).

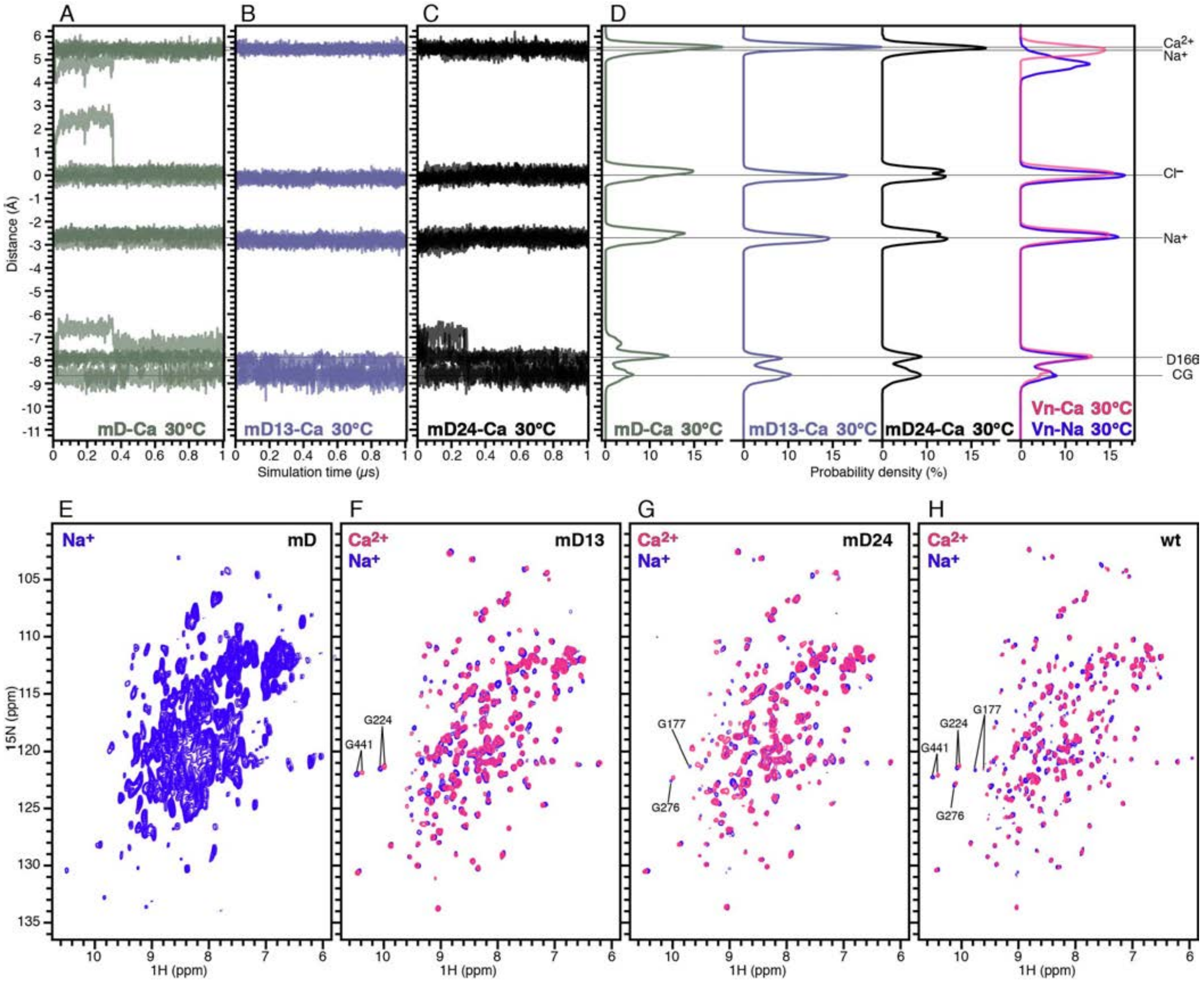

**Figure S5. MD simulations and NMR spectra of wild-type and mutant Vn HX.**

**(A-D) MD simulations of ion positions within the HX channel.** Time series (A-C) and average probability distributions (D) of the distance of $Ca^{2+}$, $Na^{+}$, and $Cl^{-}$ ions, and D166 CG atom, to the average position of the channel-occluded $Cl^{-}$ anion, for five independent MD simulations of $Ca^{2+}$-bound mutants and wild-type HX at 30°C. Ion binding sites (m1, m2, $Cl^{-}$) and D166 are marked. Each distribution is the average over the last 500 ns of five independent 1 μs MD simulations.

**(E-H)** Solution NMR 2D $^{1}H/^{15}N$ HSQC spectra of mutants mD (E), mD13 (F), mD24 (G) and wild-type (H) Vn HX, acquired without (blue) or with (pink) 2 mM $CaCl_2$.
